# Expected reward value and reward uncertainty have temporally dissociable effects on memory formation

**DOI:** 10.1101/280164

**Authors:** Jessica K. Stanek, Kathryn C. Dickerson, Kimberly S. Chiew, Nathaniel J. Clement, R. Alison Adcock

## Abstract

Anticipating rewards has been shown to enhance memory formation. While substantial evidence implicates dopamine in this behavioral effect, the precise mechanisms remain ambiguous. Because dopamine nuclei show two distinct physiological signatures of reward prediction, we hypothesized two dissociable effects on memory formation. These two signatures are a phasic dopamine response immediately following a reward cue that encodes its expected value, and a sustained, ramping dopamine response that is greater during high reward uncertainty (Fiorillo, Tobler, & Schultz, 2003). Here, we show in humans that the impact of reward anticipation on memory for an event depends on its timing relative to these physiological signatures. By manipulating reward probability (100%, 50%, or 0%) and the timing of the event to be encoded (just after the reward cue versus just before expected reward outcome), we demonstrated the predicted double dissociation: early during reward anticipation, memory formation was improved by increased expected reward value, whereas late during reward anticipation, memory formation was enhanced by reward uncertainty. Moreover, while the memory benefits of high expected reward in the early interval were consolidation-dependent, the memory benefits of high uncertainty in the later interval were not. These findings support the view that expected reward benefits memory consolidation via phasic dopamine release. The novel finding of a dissociable memory enhancement, temporally consistent with sustained anticipatory dopamine release, points toward new mechanisms of memory modulation by reward now ripe for further investigation.

## Introduction

Episodic memory formation, an important component of learning, is enhanced during reward anticipation: Just as the desire to get an ‘A’, or to understand the world, can motivate individuals to remember information, the promise of money can motivate people to form new memories (Adcock, Thangavel, Whitfield-Gabrieli, Knutson, & Gabrieli, 2006; Gruber & Otten, 2010; Wittmann et al., 2005) and even enhance memory for incidental events (Mather & Schoeke, 2011; Murty & Adcock, 2014). However, the mechanisms of memory enhancement during reward anticipation remain incompletely understood (for reviews see Miendlarzewska, Bavelier, & Schwartz, 2016; Shohamy & Adcock, 2010).

One proposed mechanism involves the neuromodulator dopamine, released during reward anticipation, which directly stabilizes long-term potentiation to support memory formation. In the dopaminergic midbrain, the ventral tegmental area (VTA) sends afferent projections to the hippocampus (Gasbarri, Sulli, & Packard, 1997; Gasbarri, Verney, Innocenzi, Campana, & Pacitti, 1994), which is populated with dopamine receptors (Bergson et al., 1995; Camps, Cortés, Gueye, Probst, & Palacios, 1989; Ciliax et al., 2000; Dawson, Gehlert, McCabe, Barnett, & Wamsley, 1986; Jiao, Paré, & Tejani-Butt, 2003; Khan et al., 2000; Lewis et al., 2001; Little, Carroll, & Cassin, 1995). Indeed, applying dopamine receptor antagonists in the hippocampus blocks memory formation for new, rewarding events (Bethus, Tse, & Morris, 2010). Prior work has also shown that during reward anticipation, activation of the dopaminergic midbrain (Adcock et al., 2006; Wittmann et al., 2005) and increased midbrain connectivity with the hippocampus (Adcock et al., 2006) predict successful memory formation. However, this mechanism of memory enhancement is only one among many known network and cellular actions of dopamine. In the hippocampus, particularly, these models must be elaborated to incorporate knowledge about dopamine receptor distributions (see Shohamy and Adcock, 2010 for review) and multiple temporal profiles of dopamine neuronal responses.

More specifically, rapid phasic burst responses scale with the expected reward value of a reward or a cue predicting reward (Fiorillo et al., 2003; Tobler, Fiorillo, & Schultz, 2005), whereas a slower, anticipatory sustained response has been reported to be associated with reward uncertainty (Fiorillo et al., 2003). Dopamine receptors in the hippocampus do not closely appose dopamine terminals (for review see Shohamy & Adcock, 2010) and the phasic responses and sustained responses are likely to differentially influence hippocampal dopamine receptors. Thus, in this study, we proposed that over several seconds of reward anticipation, phasic and sustained dopamine neuronal excitation should differentially modulate memory formation and furthermore that we could characterize these distinct dopamine profiles using a behavioral paradigm in humans. Specifically, we hypothesized that for events immediately following a reward-predicting cue, phasic dopamine release would drive memory enhancements when expected reward value is high. On the other hand, for events closer to a potentially rewarding outcome, sustained dopamine release should drive memory enhancements when reward uncertainty is high.

In addition to their association with different epochs within reward anticipation, it is unknown whether phasic and sustained dopaminergic profiles influence memory on similar or distinct timescales (i.e., requiring consolidation). Dopamine has been implicated in enhancing both early- and late-phase long term potentiation (Lemon & Manahan-Vaughan, 2006; Otmakhova & Lisman, 1996) and in increasing neuronal replay (McNamara, Tejero-Cantero, Trouche, Campo-Urriza, & Dupret, 2014). Accordingly, some of these mechanisms would have effects immediately whereas others would be apparent only after a delay (i.e., 24 hours). Thus, we also sought to establish whether putatively phasic versus sustained dopaminergic influences on memory would be present only after a period that allowed for consolidation, or would be evident immediately after encoding.

We set out to dissociate the putative influence of two distinct dopaminergic responses on memory formation during reward anticipation. To parse these effects, we designed a study in which we used overlearned abstract cues to indicate reward probability, establishing expected reward value independently from uncertainty. We further manipulated the epoch of encoding during reward anticipation: we presented items either early (400ms after cue presentation), to capture a rapid dopamine response anticipated to scale with expected reward value, or late (3-3.6s after cue presentation), to capture a sustained dopamine response anticipated to scale with high reward uncertainty. Finally, we manipulated retrieval time, either 15 minutes- or 24 hours-post encoding, to examine the effects on memory performance with and without consolidation. With this paradigm, therefore, we examined how expected reward value and reward uncertainty each influenced memory formation.

## Methods

### Subjects

Forty healthy young adult volunteers participated in the study. All participants provided informed consent, as approved by the Duke University Institutional Review Board. Data from additional participants were excluded due to failure to follow the instructions (n =1), poor cue-outcome learning (n = 2) or computer error (n = 3). Individuals participated in one of two experiments: Experiment 1 (n = 20, 12 female, mean age = 27.45 ± 3.82 years) or Experiment 2 (n = 20, 12 female, mean age = 21.90 ± 3.23 years).

### Design and Procedure

#### Reward Learning

The first phase of the experiment involved reward learning. During reward learning, participants were presented with abstract cues, all Tibetan characters, which predicted 100%, 50%, or 0% probability of subsequent monetary reward. Participants were instructed to try to learn the relationship between the cues and reward. They were presented with the cue (1s), a unique image of an everyday object (2s), then an image of either a dollar bill or a scrambled dollar bill (400ms), indicating a reward or no reward respectively. A jittered fixation-cross separated trials (1-8s). No motor contingency was required to earn the reward. Independent of performance, participants were paid a monetary bonus equal to the amount accumulated over the outcomes in one block of the task. Participants saw 40 trials per condition, distributed evenly over five blocks. Prior to the first block and following every block, participants were asked to rate their certainty of receiving reward following each cue along a sliding scale from “Certain: No Reward” to “Certain: Reward.” To be included in the analysis, during learning participants had to meet a minimum criteria of identifying the 100% reward cues as more associated with reward than the 0% cues, as assessed by average certainty score across all 5 blocks.

#### Incidental Encoding

In the second phase of the experiment, the abstract cues used in the reward learning phase were used to modulate incidental encoding; these cues predicted 100%, 50%, and 0% reward probability. Because the associations were deterministic, reward probabilities established expected reward value, with 100% higher than 50% and 0% rewarded cues. In contrast, 50% predictive cues established higher uncertainty relative to the 100% and 0% predictive cues.

During the incidental encoding task, participants saw a cue (400ms), followed by a unique novel object (1s) either immediately after the cue (400ms post-cue onset and 3.2s pre-outcome) or just prior to outcome (3-3.6s post-cue onset and 0-0.6s pre-outcome). These encoding epochs were chosen based on the timing of the phasic dopamine response [<500ms (Schultz, Dayan, & Montague, 1997)] and the sustained ramping response [2s (Fiorillo et al., 2003); also 4-6s (Howe, Tierney, Sandberg, Phillips, & Graybiel, 2013; Totah, Kim, & Moghaddam, 2013)]. When no image was present during the delay, a fixation cross was shown. A dollar bill or scrambled dollar bill, indicating a reward or no reward respectively (400ms), appeared 3.6 s after cue onset for all trials. After reward feedback, participants were presented with the probe question, “Did you receive a reward?” (1s). Participants were instructed to quickly and accurately make a “yes” or “no” button press. The exact motor component could not be anticipated since the yes/no, right/left location was random from trial-to-trial. A jittered fixation-cross separated trials (1-7s). In sum, there were six conditions in the design: three probabilities of reward (100%: high expected reward value/certain, 50%: medium expected reward value/uncertain, and 0%: no expected reward value/certain) crossed with the early or late encoding epochs (**Figure 1**). There were 20 trials per condition, evenly dispersed among five blocks.

**Figure 1.**
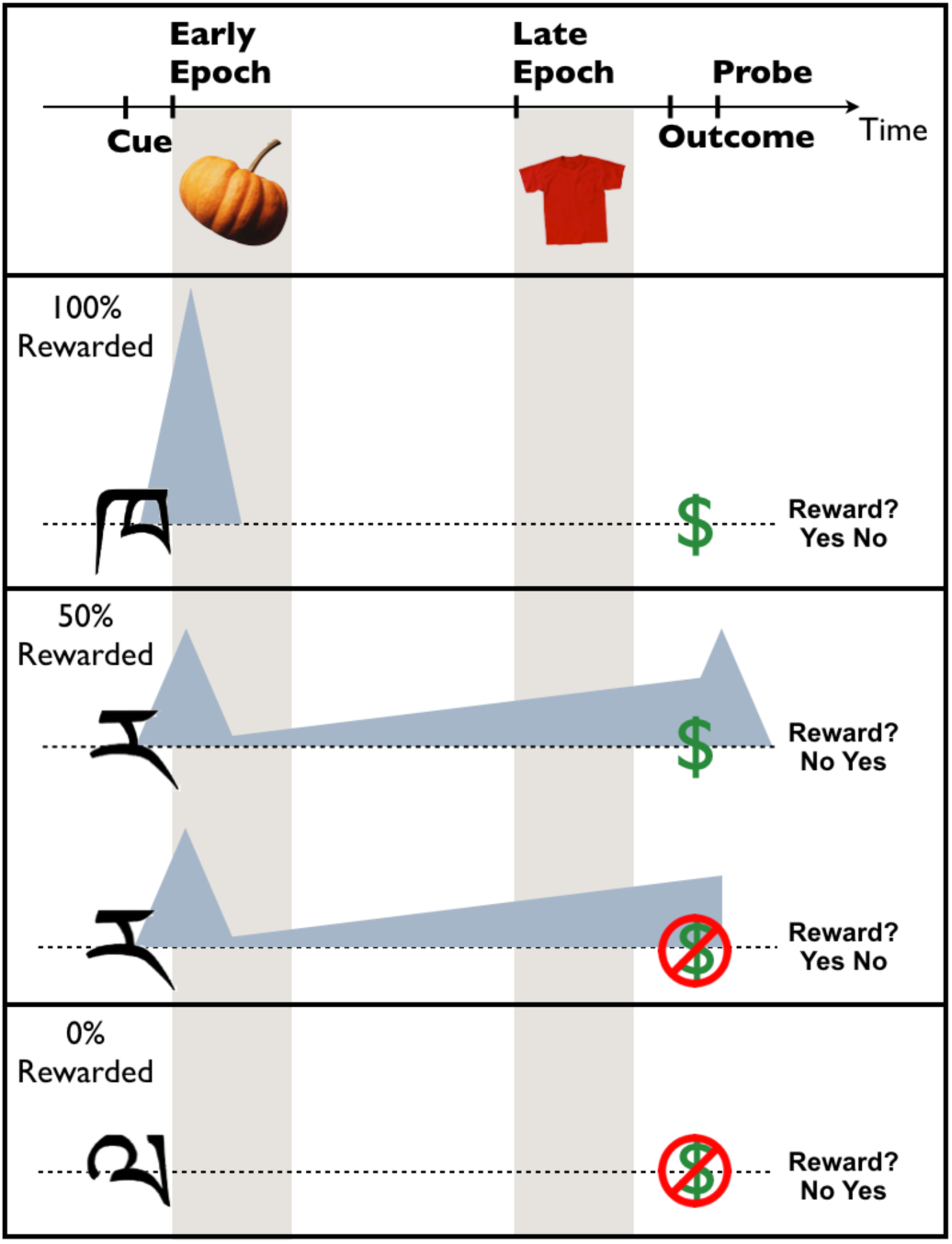
Experimental Design: Incidental Encoding Task. The task was designed to dissociate two physiological profiles of a putative dopamine response during reward anticipation - a phasic response that occurs rapidly and scales with expected reward value (Early Epoch) and a sustained response that increases with uncertainty (Late Epoch). Shaded triangles indicate these dopamine profiles relative to the cue, encoding epochs, outcome, and reward probe events in each trial. Cues associated with 100%, 50%, or 0% reward probability were presented for 400ms. Incidental encoding objects were presented for 1s either immediately following the cue (400ms post-cue onset) or shortly before anticipated reward outcome (3.2s pre-reward delivery). Finally, a probe asking participants whether or not they received a reward, with the yes/no response mapping counterbalanced, followed for 1s.

#### Recognition Memory Test

Participants performed an old/new recognition memory test where they viewed 280 “new” objects and 280 “old” objects. Although “old” objects were from both the reward learning and incidental encoding phases, only the old objects from the incidental encoding phase were included in analyses to calculate memory performance. They rated their confidence by saying “Definitely Sure,” “Pretty Sure,” or “Just Guessing” for each memory judgment. One-sample t-tests examined whether memory for guesses (within guesses: [hits – false alarms]/all responses) was above chance in each condition. Significant memory for guesses resulted in trials of all confidence being included in the analysis for both groups.

#### Experiment 1 – 24-Hour Retrieval

In Experiment 1, participants returned at the same time the next day to complete the recognition memory test, approximately 24 hours after encoding.

#### Experiment 2 – Immediate Retrieval

In Experiment 2, participants completed the recognition memory test 15 minutes after completing the encoding task.

### Analysis

Memory performance for all analyses was calculated as a corrected hit rate ([hits – false alarms]/all responses).

#### Within Experiments

A 3 × 2 repeated-measures ANOVA was used to examine the effects of reward probability (100%, 50%, 0%) and encoding epoch (early, late) on subsequent memory performance. Any significant interaction between reward probability and encoding epoch warranted post-hoc analyses. One-way repeated-measures ANOVAs at early encoding (400ms post-cue onset) and late encoding (3.2s pre-reward delivery) were used to examine how reward probability related to memory formation at each encoding epoch during anticipation. Significant one-way ANOVAs prompted follow-up analyses. Specifically, a test for a linear trend increasing with probability was used to examine how expected reward value related to memory, and paired Student’s t-tests were used to compare memory on certain (100%, 0%) versus uncertain (50%) trials.

To determine whether memory for items presented following the 50% probability cue was influenced by reward outcome, we completed a two-tailed paired Student’s t-test to see if there were differences in memory for rewarded versus unrewarded trials within that condition. These were conducted separately at the early and late encoding epochs.

To examine whether variability in attention and task engagement at encoding could account for subsequent memory performance across conditions, we examined performance on the reward probe; we conducted 1-way ANOVAs and follow-up pairwise Student’s t-tests and tests for a linear trend to determine whether reaction time or accuracy for the reward probe varied by condition.

#### Across Experiments

Since both Experiment 1 and Experiment 2 revealed differences in the pattern of memory formation for early versus late encoding epochs, we tested whether the patterns at each encoding epoch significantly differed according to retrieval time. We thus performed 3 × 2 ANOVAs with reward probability (100%, 50%, 0%) as a within-subjects factor and retrieval time (immediate, 24-hours) as an across-subjects factor. We conducted this analysis at both the early and late encoding epochs. A significant interaction between reward probability and retrieval time prompted post-hoc pairwise ANOVAs to examine whether the deltas between immediate and 24-hour retrieval were significantly different across reward probability conditions.

## Results

### Reward Learning

Participants in both groups successfully learned the meaning of the cues during the reward-learning phase. In the 24-hour memory group, participants in the final block reported the 100% probable cue as 99.46% (± 0.20 SEM) likely to predict reward, the 50% cue as 52.65% (± 3.39 SEM) likely to predict reward and the 0% cue as 2.29% (± 1.79 SEM) likely to predict reward. In the immediate memory group, participants in the final block reported the 100% probable cue as 99.37% (± 0.43 SEM) likely to predict reward, the 50% cue as 55.26% (± 2.68 SEM) likely to predict reward and the 0% cue as 1.44% (± 1.07 SEM) likely to predict reward.

### Experiment 1 – 24-Hour Retrieval

Because the aim of the study was to manipulate distinct temporal components of reward anticipation and relate those components to determinants of dopamine physiology, we completed a 3 × 2 ANOVA looking at memory performance as a function of reward probability and encoding epoch. We found a main effect of reward probability (F(2,18) = 5.56, p = 0.01), no main effect of encoding epoch (F(1,19) = 2.57, p = 0.13), and a strong interaction between encoding epoch and reward probability (F(2,18) = 7.50, p = 0.004). Follow-up one-way ANOVAs examining the effect of reward probability on memory within early and late encoding epochs revealed a significant effect in the late epoch (F(2,19) = 13.25, p < 0.0001) and a trend-level effect in the early epoch (F(2,19) = 2.41, p = 0.10).

Post-hoc tests to examine memory performance during the early encoding epoch revealed a significant linear trend such that memory scaled with increasing reward probability (Linear trend: R square = 0.03, p = 0.04). Thus, early during anticipation, memory performance linearly tracked expected reward value (**Figure 2A**).

**Figure 2.**
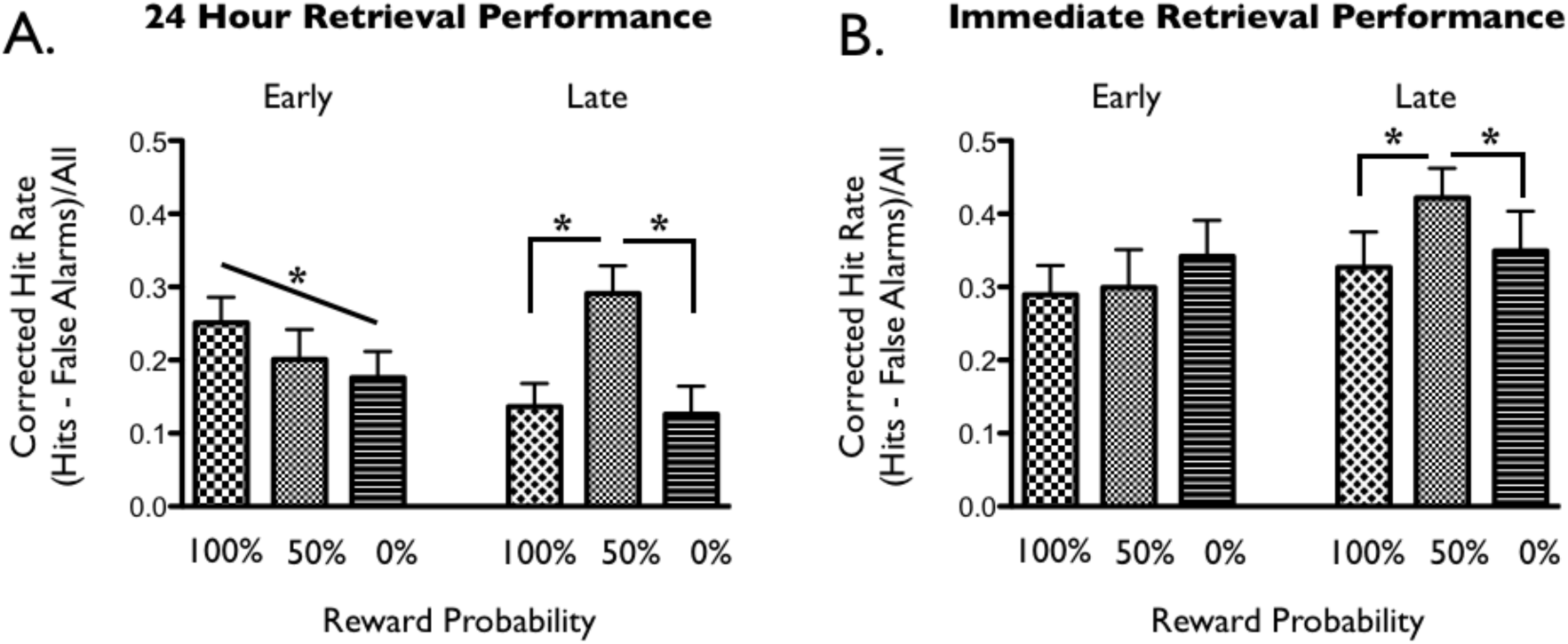
Memory Performance for Early and Late Encoding Epoch at 24-Hour and Immediate Retrieval. **A.** In the 24-hour Retrieval group, Early Epoch memory linearly increased with expected reward value; Late Epoch memory was greatest for items encoded during reward uncertainty. **B.** In the Immediate Retrieval group, Early Epoch memory did not differ by expected reward value. As in the 24-hour Retrieval group, Late Epoch memory for the Immediate Retrieval group was greatest for items encoded during reward uncertainty.

Post-hoc pairwise t-tests to examine memory performance during the late encoding epoch revealed greater memory following 50% cues vs. either 100% or 0% cues, with no difference in performance between 100% and 0% cues (100% vs. 50%: t(19) = 4.34, p = 0.0004; 50% vs. 0%: t(19) = 4.20, p = 0.0005; 100% vs. 0%: t(19) = 0.31, p = 0.76). Thus, late in reward anticipation, greater reward uncertainty benefitted memory (**Figure 2A**).

It was also possible that the memory benefit we attributed to the uncertain anticipatory context could instead be explained by associations with reward outcomes. To investigate this alternative explanation, we performed t-tests between the rewarded and unrewarded uncertain trials, during both early and late epochs. We found no differences in memory as a function of reward outcome in either epoch (early, rewarded vs. unrewarded: t(19) = 0.10, p =0.92; late, rewarded vs. unrewarded: t(19) = 1.09, p = 0.29).

### Experiment 2 – Immediate Retrieval

The 24-hour retrieval test did not allow us to distinguish between effects acting at encoding versus consolidation. Thus, in Experiment 2, participants completed an immediate retrieval test, 15 minutes after encoding. All analyses for Experiment 1 were repeated for Experiment 2. Analyses of immediate retrieval performance replicated effects of reward uncertainty on items presented late in the anticipation epoch; however, they did not show effects of reward probability on items presented early in the epoch, as follows:

A 3 × 2 ANOVA revealed a trend for a main effect of reward probability (F(18) = 3.33, p = 0.06) and a main effect of encoding epoch (F(19) = 8.008, p= 0.01), with memory greater at late than early encoding epochs. Importantly, there was again an interaction between reward probability and encoding epoch (F(18) = 3.711, p = 0.04). Post-hoc one-way ANOVAs within early and late encoding epochs revealed a significant difference in memory for the late epoch (F(2,19) = 4.95, p = 0.01) but no difference for the early epoch (F(2,19) = 1.31, p = 0.28).

Replicating the memory benefit observed in the 24-hour retrieval condition for reward uncertainty in the late encoding epoch, follow-up t-tests again revealed a difference following 50% cues relative to both 100% and 0% cues, with no difference between the latter (100% vs. 50%: t(19) = 2.97, p = 0.008; 50% vs. 0%: t(19) = 2.57, p = 0.02; 100% vs. 0%: t(19) = 0.66, p = 0.52), The presence of the uncertainty effect at both immediate and 24-hour retrieval indicates that this effect was not dependent on consolidation (**Figure 2B**).

By contrast, the effect of expected reward value for items in the early encoding epoch was not present at immediate retrieval. Although the ANOVA demonstrated no significant difference by reward probability, in Experiment 2, the test for a linear trend was an a priori analysis. We found no significant linear trend (Rsquare = 0.01, p = 0.14). Thus, the influence of reward probability on memory for items presented early during reward anticipation was not present during immediate retrieval, and only appeared after 24 hours (**Figure 2B**).

As was the case at 24-hour retrieval, analyses for immediate retrieval revealed no effects of reward outcome on memory during uncertain trials (early, rewarded vs. unrewarded t(19) < 0.0001, p = 1.00; late, rewarded vs. unrewarded t(19) = 1.33, p = 0.20).

### Experiments 1 & 2: Contrasting 24-Hour and Immediate Memory Results

To quantify whether memory patterns within early and late encoding epochs changed over a 24-hour period of consolidation, we ran two 3 × 2 ANOVAs, one per encoding epoch, with the factors reward probability and retrieval time and looked for an interaction between the two. We found a significant interaction at the early epoch (F(2,37) = 4.281, p = 0.021) but not at the late epoch (F(2,37) = 1.826, p = 0.175). Follow-up pairwise ANOVAs revealed that the decrement in memory performance as a function of retrieval period (24-hour vs. immediate) was significantly greater for 0% than 100% reward (F(1,38) = 8.76, p = 0.005), with no other significant differences (all other ANOVAs: F(1,38) < 1.75, p > 0.19). After consolidation, memory for items encoded during the early epoch decreased more following 0% cues than following 100% cues. Thus, the relationship between memory and reward anticipation remained consistent from immediate to 24-hour retrieval for items at the late encoding epoch, but changed significantly across retrieval periods in the early encoding epoch (**Figure 3**).

**Figure 3.**
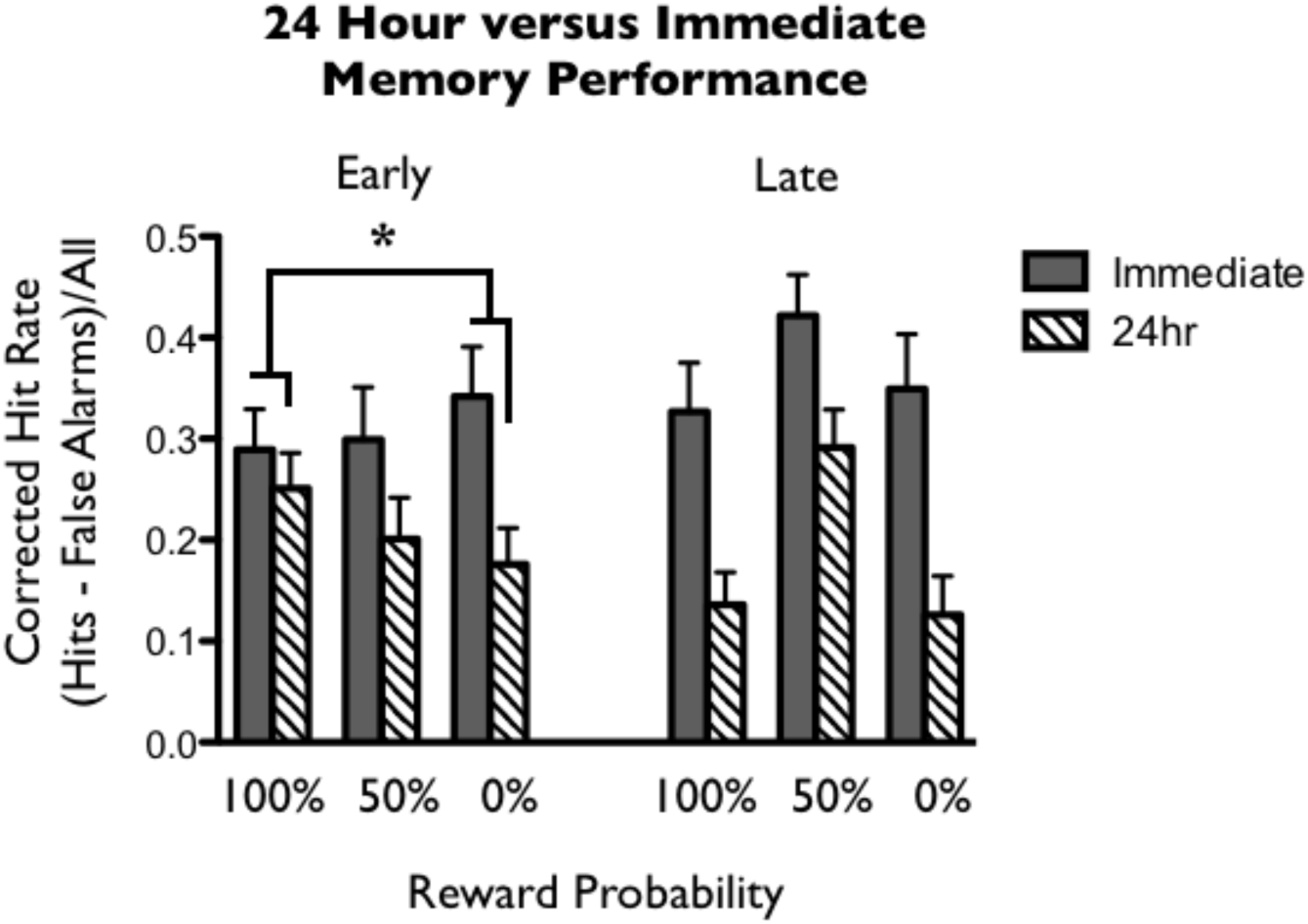
Retrieval Group by Expected Reward Value Interaction. Within the Early Epoch, there was a significant interaction between reward probability (100%, 50%, 0%) and retrieval group (Immediate, 24-hour). The difference between the 24-hour and Immediate Retrieval groups was smaller at 100% reward probability and greater at 0% reward probability. Thus in the Early Epoch, the impact of reward value on memory is only evident after overnight consolidation.

### Experiments 1 & 2: Accuracy and Reaction Time During Encoding

To examine potential contributions of task engagement at encoding to the observed relationships between reward anticipation and memory, we examined whether the patterns of accuracy or reaction time for reward probes resembled subsequent memory performance across conditions. In both retrieval groups, 1-way ANOVAs showed no accuracy differences in probe response by reward probability for trials with items presented in the early epoch (24-hour: F(2,19) = 2.37, p = 0.11; Immediate: F(2,19) = 1.42, p = 0.25; **Figure 4A and 4C**). However, probe accuracy on trials with items presented in the late epoch in both groups significantly differed with reward probability (24-hour: F(2,19) = 9.83, p = 0.0004; Immediate: F(2,19) = 9.58, p < 0.0001), and revealed significant linear trends, such that people performed more accurately as expected reward value increased (24-hour: R square = 0.06, p < 0.0001; Immediate: R square = 0.04, p = 0.004; **Figure 4A and 4C**). One-way ANOVAs of reaction time on trials with items presented in the late epoch revealed a significant difference at 24-hour but not immediate retrieval (24-hour: F(2,19) = 4.25, p = 0.02; Immediate: F(2,19) = 1.97, p = 0.15; **Figure 4B and 4D**).

**Figure 4.**
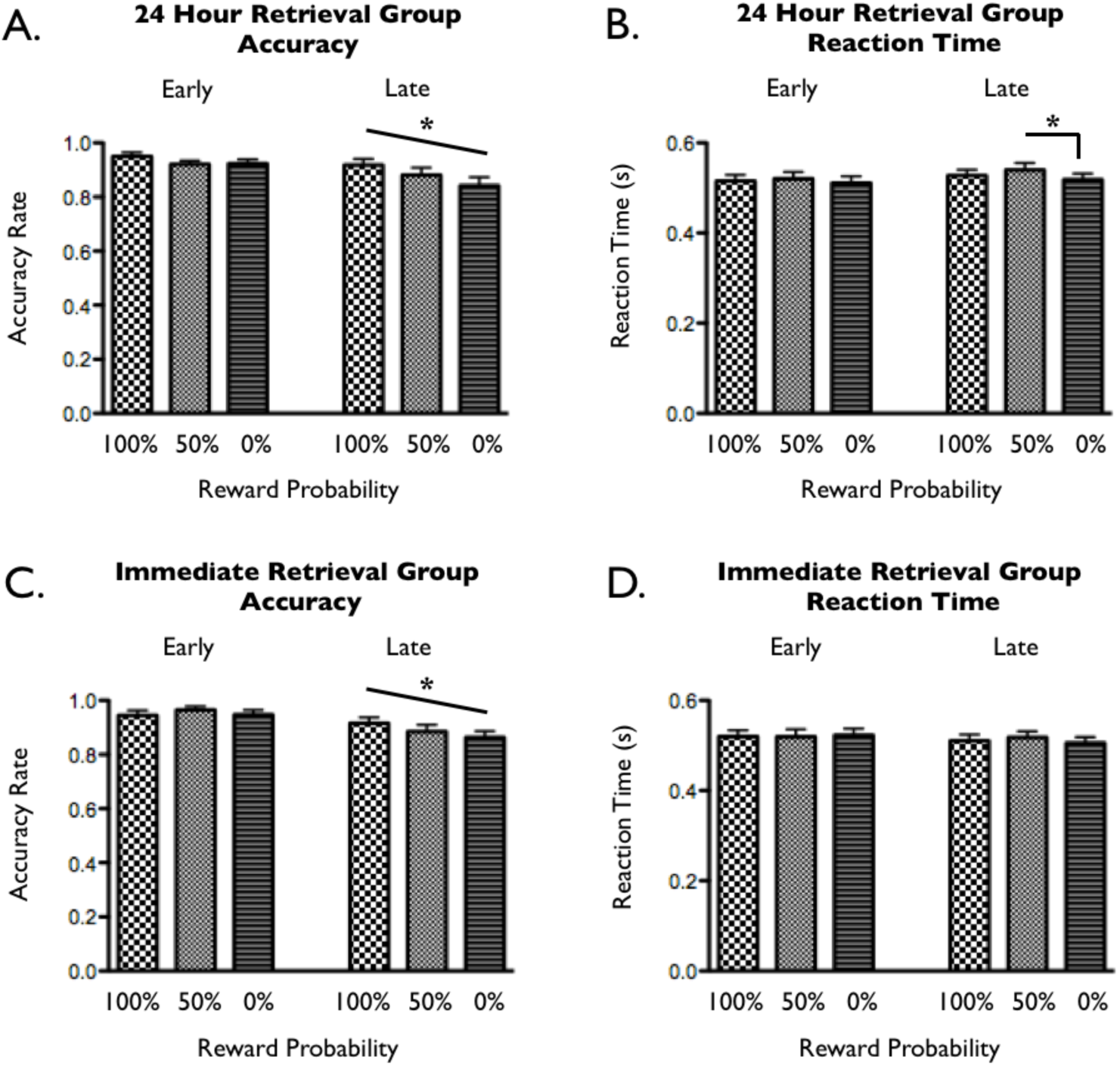
Accuracy and Reaction Times for Reward Probe at Encoding. **24-hour retrieval group: A.** Unlike memory, accuracy for responding to the yes/no reward probe on trials with items presented in the early epoch did not differ by expected reward value. For trials with items in the late epoch, accuracy linearly increased with increasing expected reward value. **B.** Reaction times on trials with items presented in the early epoch did not differ by expected reward value. Reaction times on trials with items presented in the late epoch were slower for the 50% trials than the 0% trials. **Immediate retrieval group: C.** The pattern of accuracy for detecting the yes/no reward probe was similar to the 24-hour retrieval group. **D.** There were no significant differences in reaction time for any condition.

Follow-up pairwise t-tests showed that the 24-hour effects were driven by slower reaction times for 50% rewarded trials relative to 0% rewarded trials (50% vs. 0%: t(1,19) = 3.84, p = 0.001; 100% vs. 50%: t(1,19) = 1.44, p = 0.17, 100% vs. 0%: t(1,19) = 1.18, p = 0.25; **Figure 4B**). There were no significant reaction time differences on trials with items presented in the early epoch (24-hour: F(2,19) = 1.12, p = 0.24; Immediate: F(2,19) = 0.15, p = 0.86; **Figure 4B and 4D**). Thus, reaction time and accuracy were modulated by reward probability, but not in a manner that paralleled the observed memory effects.

## Discussion

Our findings demonstrate temporally specific reward anticipation influences on memory formation. During an early encoding epoch, 400ms after the presentation of the reward cue, and temporally coincident with phasic dopamine responses, item memory scaled with expected reward value. During a late encoding epoch, just prior to a predicted outcome, memory was instead greatest for items presented during high uncertainty. The uncertainty benefit was present both immediately and 24 hours after encoding, implying an underlying mechanism modulating memory formation at encoding. The memory benefit for expected reward value, however, emerged only after 24 hours, implying a distinct mechanism acting during consolidation to modulate memory formation.

### Dopaminergic Accounts

While this is, to our knowledge, the first behavioral demonstration of dissociable temporal contexts for encoding within reward anticipation, the results build on expectations generated from prior neuroimaging and physiological studies. Previous work using functional magnetic resonance imaging (fMRI) has demonstrated dissociable neural responses within the dopaminergic system for expected reward value and uncertainty (Preuschoff, Bossaerts, & Quartz, 2006; Tobler, O’Doherty, Dolan, & Schultz, 2007), with one study demonstrating dissociable temporal patterns in the striatum: activation in the first second after a reward cue scaling with expected reward value and in the following seconds leading up to reward outcome scaling with uncertainty (Preuschoff et al., 2006). Physiologically, cues associated with greater expected reward value elicit greater phasic dopamine firing in the midbrain at latencies less than 400ms (Fiorillo et al., 2003; Tobler et al., 2005), whereas a sustained dopaminergic ramp has been shown to increase with greater reward uncertainty over a two second period of reward anticipation (Fiorillo et al., 2003). Because phasic dopamine would be predicted to benefit memory early in the reward anticipation period and sustained anticipatory dopamine to have an effect close to the reward outcome, the present study implies previously undescribed, functionally specific relationships between memory and phasic versus sustained dopamine.

Dissociable effects on memory are grounded in observations of other differential effects of dopaminergic firing modes. Phasic burst firing preferentially influences downstream targets via synaptic release, while sustained low-frequency activity results in in extrasynaptic release (Floresco, West, Ash, Moore, & Grace, 2003). Extracellular dopamine levels have been demonstrated not only to increase for increased tonic dopamine activity (Floresco et al., 2003), but also to exhibit sustained, ramping dopamine levels lasting on the order of seconds (Howe et al., 2013; Roitman, Stuber, Phillips, Wightman, & Carelli, 2004; Stuber, Roitman, Phillips, Carelli, & Wightman, 2005). The mismatch of distribution of dopamine receptors in the hippocampus relative to dopamine terminals indicates that phasic dopamine firing in the midbrain cannot be communicated to hippocampal synapses as a temporally precise signal (see Shohamy & Adcock 2010 for review). Optogenetic findings have also revealed that higher (simulating phasic) versus lower (simulating tonic, or possibly sustained) levels of dopamine release have differential influences on dopamine receptors in the hippocampus (Rosen, Cheung, & Siegelbaum, 2015). While further work remains to elucidate phasic versus sustained (or tonic) dopamine effects on memory, the extant literature supports multiple dissociable mechanisms of dopaminergic influence on the hippocampus at distinct timescales (Düzel, Bunzeck, Guitart-Masip, & Düzel, 2010; Shohamy & Adcock, 2010).

The present observation of a 24-hour memory benefit following higher expected reward value early during reward anticipation is consistent with a previously described relationship in the literature between dopamine and consolidation-dependent memory effects. Expected reward value predicts phasic dopamine activity in the VTA (Cohen, Haesler, Vong, Lowell, & Uchida, 2012; Fiorillo et al., 2003; Pan, Schmidt, Wickens, & Hyland, 2005; Schultz, 1998; Schultz et al., 1997; Tobler et al., 2005). Dopamine has been associated with enhancement of late-phase long term potentiation (Lemon & Manahan-Vaughan, 2006) and is a critical element in the synaptic-tagging and capture theory of memory consolidation (Lisman, Grace, & Duzel, 2011; Redondo & Morris, 2011; Sajikumar & Frey, 2004). Additionally, optogenetically-induced burst firing of dopaminergic fibers results in increased hippocampal replay during sleep and increased memory (McNamara et al., 2014). Thus, previous work supports a relationship between dopamine and enhanced hippocampal memory consolidation. Our results during the early epoch of reward anticipation, a novel demonstration of a temporally-specific effect of expected reward value on memory, are consistent with phasic dopamine driving consolidation-dependent memory processes.

On the other hand, our observation of an immediate memory benefit just prior to reward outcome during high uncertainty suggests a mechanism of memory enhancement that occurs at encoding, independent of consolidation. High reward uncertainty has been associated with sustained, ramping dopamine firing in the VTA (Fiorillo et al., 2003). While our results are consistent with a modulatory influence of sustained dopamine release on memory, a relationship between sustained dopamine release and hippocampal memory formation has yet to be demonstrated. What has been shown, however, is that tonic dopamine has immediately observable effects on hippocampal physiology (Rosen et al., 2015) and early long term potentiation (Li, Cullen, Anwyl, & Rowan, 2003), providing candidate mechanisms by which sustained dopamine may contribute to hippocampal memory at immediate retrieval. This hypothesis opens new avenues for future investigation.

By demonstrating and dissociating both immediate and consolidation-dependent memory benefits related to reward anticipation, our study takes an important first step towards reconciling conflicting patterns of findings in the memory literature. Prior rodent work has demonstrated the importance of consolidation for dopamine-dependent memory formation (Bethus et al., 2010; McNamara et al., 2014) and theoretical mechanisms of dopamine synaptic activity have emphasized consolidation (Lisman et al., 2011; Redondo & Morris, 2011). However, some reward anticipation effects on memory are not consolidation-dependent (Gruber, Gelman, & Ranganath, 2014; Murty & Adcock, 2014). Our data integrates across previous studies, suggesting that phasic, synaptic dopamine effects may be reliant on consolidation, whereas sustained, extrasynaptic dopamine effects may occur at encoding.

### Alternative Accounts

In the present study, we did not manipulate dopamine directly. It is thus possible that our effects, and in particular, the uncertainty benefit modulating encoding, were not dopaminergic in nature. Although prior studies using pharmacological manipulations have already contributed direct evidence that dopamine affects memory formation in humans (Chowdhury, Guitart-Masip, Bunzeck, Dolan, & Düzel, 2012; Knecht et al., 2004), they have not been shown to selectively affect specific modes of dopamine firing. Our hypotheses were based on work showing sustained neuronal firing in the VTA scaling with greater uncertainty. It has been debated whether this signal represents sustained dopaminergic firing or represents an accumulation of phasic responses (Niv, Duff, & Dayan, 2005). Other work, however, is consistent with a sustained signal that is actively maintained (Howe et al., 2013; Lloyd & Dayan, 2015; MacInnes, Dickerson, Chen, & Adcock, 2016; Murty, Ballard, & Adcock, 2016; Totah et al., 2013). There is evidence, on the other hand, that ramping activity in the VTA may be GABAergic in nature (Cohen et al., 2012). In addition, sustained dopamine release in efferent regions has been demonstrated scaling with reward proximity and reward magnitude (Howe et al., 2013); a response to uncertainty has yet to be experimentally examined in efferent regions. Other neurotransmitters, such as acetylcholine, offer additional potential mechanisms for enhanced memory formation that are not mutually exclusive. The hippocampus is densely populated with cholinergic receptors (Alkondon & Albuquerque, 1993) and acetylcholine has been discussed as important for expected uncertainty (Sarter, Lustig, Howe, Gritton, & Berry, 2014; Yu & Dayan, 2005), which may be similar to the cued uncertainty in this study. Finally, under some behavioral contexts, dopamine release in the hippocampus appears to require neuronal activity within the locus coeruleus, implicating by noradrenergic neurons (Kempadoo, Mosharov, Choi, Sulzer, & Kandel, 2016; Smith & Greene, 2012; Takeuchi et al., 2016). Thus, the current work introduces possibilities for future experiments that disentangle the roles of specific neuromodulators in encoding during high reward uncertainty.

Multiple alternative accounts were also considered as potential explanations of our memory findings. One intuitive possibility is that enhanced memory formation was a result of greater task engagement; however, this alternative account was not supported by reaction time nor accuracy data. Another possible alternative was that the memory benefit for uncertainty in the late encoding epoch was due to a phasic dopaminergic response to reward delivery. However, there was no evidence for this relationship: there was no memory difference for items presented prior to rewarded versus unrewarded outcomes on the uncertain trials.

### Conclusion

The present study builds on prior findings that reward anticipation modulates memory formation to show that within reward anticipation, there are distinct temporal contexts for encoding, with mechanistically distinct impact on memory outcomes. Moreover, by mapping these encoding contexts onto the putative physiological profiles for expected reward value and uncertainty, this work suggests a novel and testable model of dopaminergic influence on memory formation: that sustained dopamine release acts to benefit memory encoding, whereas phasic dopamine release acts to facilitate memory consolidation. Integrating disparate findings, our proposed model paves the way for future research examining contextually-regulated mechanisms of reward-enhanced memory formation.

## Acknowledgements

The authors would like to thank Vishnu Murty for helpful comments on this manuscript. This work was supported by a NIMH BRAINS award (R01MH094743) to R.A.A and a KL2 award (UL1TR001117) to K.C.D. The authors declare no competing financial interests.

